# Exosome-like biogenesis from the Golgi releases extracellular vesicles lacking conventional tetraspanins that mediate immune evasion in cancer

**DOI:** 10.64898/2026.05.13.719612

**Authors:** Ikjot S. Sohal, Sydney N. Shaw, Andrea P. dos Santos, Bennett D. Elzey, Haley Anne Harper, Carli McMahan, Lauren N. Meeks, Humna Hasan, Zulaida Soto-Vargas, Noor Abdullah, Subhransu Sekhar Sahoo, Majid Kazemian, Matthew R. Olson, Andrea L. Kasinski

## Abstract

Immunotherapy has improved survival across multiple malignancies but remains largely ineffective in solid cancers such as lung, breast, and pancreatic cancer. A key driver of resistance is the immunosuppressive tumor microenvironment (TME). Although numerous mediators of TME immunosuppression have been identified, therapeutic targeting has provided limited clinical benefit. Tumor-derived extracellular vesicles (EVs) have recently emerged as contributors to resistance, yet their mechanisms remain unclear. We developed human non-small cell lung cancer models to investigate EV-mediated immunosuppression. We identified a distinct Golgi-derived EV subpopulation that potently suppress T cell function and tumor infiltration. These EVs express the *trans*-Golgi network marker TGOLN2, and exhibit minimal levels of canonical EV markers. TGOLN2 overexpression drives this suppressive phenotype. Clinically, elevated *TGOLN2* associates with poor survival and correlate with an immunosuppressive TME signature across more than 20 cancer types, including NSCLC. Collectively, this work defines a previously unrecognized mechanism of TGOLN2-driven, EV-mediated immunosuppression.

**Statement of Significance:** We revealed TGOLN2 overexpression as a new immune evasion mechanism that mediates T cell suppression through increased secretion of a Golgi-derived extracellular vesicle (EV) subpopulation. These findings redefine current paradigms of EV biology and nominate TGOLN2 as a potential biomarker and therapeutic target in immunosuppressive cancers.

## INTRODUCTION

Lung and bronchus cancer are the leading cause of cancer-related deaths worldwide^1^. In the United States, it is estimated that >100,000 people will die from lung cancer in 2026^2^. Non-small cell lung cancer (NSCLC) is the most common type of lung cancer, accounting for 84% of all diagnosed cases. Therapeutically, tyrosine kinase–based targeted therapies are the most common; nonetheless, acquired resistance to these therapies evolves, leading to rapid clinical relapse and disease progression^3^. Checkpoint immunotherapies have resulted in some success in NSCLC; still, poor clinical efficacy is observed for 50% of patients treated with anti-PD1 or anti-CTLA4^4^. The immunosuppressive tumor microenvironment (TME), hallmarked by a prominent myeloid presence, in combination with poor infiltration and inactivation of T cells, is a major contributor to treatment failure^5^. Indeed, infiltration of T cells in the tumor has already been confirmed to be of positive prognostic value across 17 cancer types, including NSCLC^5^. Mechanisms proposed to mediate the immunosuppressive TME include tumor-intrinsic factors such as lack of tumor-specific antigens or c-MET amplification^6^, and tumor-extrinsic factors such as defective recruitment and activation of antigen-presenting cells^7^ or presence of immunosuppressive stroma^8^. However, current therapeutics that target these mechanisms have shown modest clinical benefit, indicating that other factors contribute to this process. One emerging mechanism involves cancer-derived extracellular vesicles (EVs), which have been shown to carry immunosuppressive capabilities and inhibit the efficacy of immunotherapy^9^.

EVs are a heterogenous population of membrane-bound vesicles secreted nearly all cell types. Recognized subtypes include exosomes, microvesicles, apoptotic vesicles, and more recently described novel subtypes^10,11^. Classification, particularly of exosomes, has traditionally relied on the presence of associated tetraspanin proteins, most notably CD63, CD9, and CD81. Owing to their ubiquitous distribution, capacity to traverse biological barriers, and dynamic regulation in response to pathophysiological changes, EVs serve not only as promising biomarkers but also as active mediators of disease progression including cancer^12^. Tumor-derived EVs have been shown to contribute to several cancer hallmarks, including metabolic reprogramming, extracellular matrix remodeling, angiogenesis, immune evasion, and metastasis^13^. However, these functions are attributed to the heterogenous population of EVs released by the cancer cells or the tumor model, leaving a significant gap in our understanding of the specific contributions of different EV subpopulations and EV biogenesis mechanisms driving these processes.

Here, using NSCLC as a model, we identified a specific subpopulation of cancer-derived EVs that strongly suppress T cell proliferation and reduces T cell tumor infiltration. We determined that the T cell-suppressive EVs originated from the Golgi and identified overexpression of TGOLN2, a classic *trans*-Golgi network (TGN) marker, as the underlying mechanism driving EV-mediated T cell suppression. In another NSCLC model that had no effect on T cells, overexpressing TGOLN2 was sufficient to increase T cell-suppressive EV release and enhance T cell suppression, validating these findings. Analysis of The Cancer Genome Atlas (TCGA) patient data supported these findings, where higher *TGOLN2* transcript levels negatively correlated with patient survival in lung cancer and other cancer types. In a pan-cancer analysis, higher *TGOLN2* levels significantly correlated with an immunosuppressive TME signature – reduced infiltration of CD4+ and CD8+ T cells and NK cells, and increased infiltration of M2 macrophages. This correlation of TGOLN2 with an immunosuppressive TME signature was observed in over 20 cancer types, including NSCLC. Collectively, these findings support a role for TGOLN2 and TGOLN2+ EVs as important mediators of immunosuppression in the TME.

## RESULTS

### Cancer-derived EVs from a subset of NSCLC cells selectively suppress T cell function

To determine whether tumor-derived extracellular vesicles (EVs) directly contribute to T cell dysfunction, we evaluated EVs derived from multiple non-small cell lung cancer (NSCLC) models for their ability to modulate T cell response. EVs isolated from H358 cells robustly suppressed Jurkat T cell proliferation, accompanied by increased necrosis and diminished T cell activation (**Fig 1a-b**). In contrast, EVs derived from A549 cells had no measurable impact on T cell proliferation, identifying a clear functional divergence between NSCLC-derived EV populations (**Fig 1a**).

**Figure 1.**
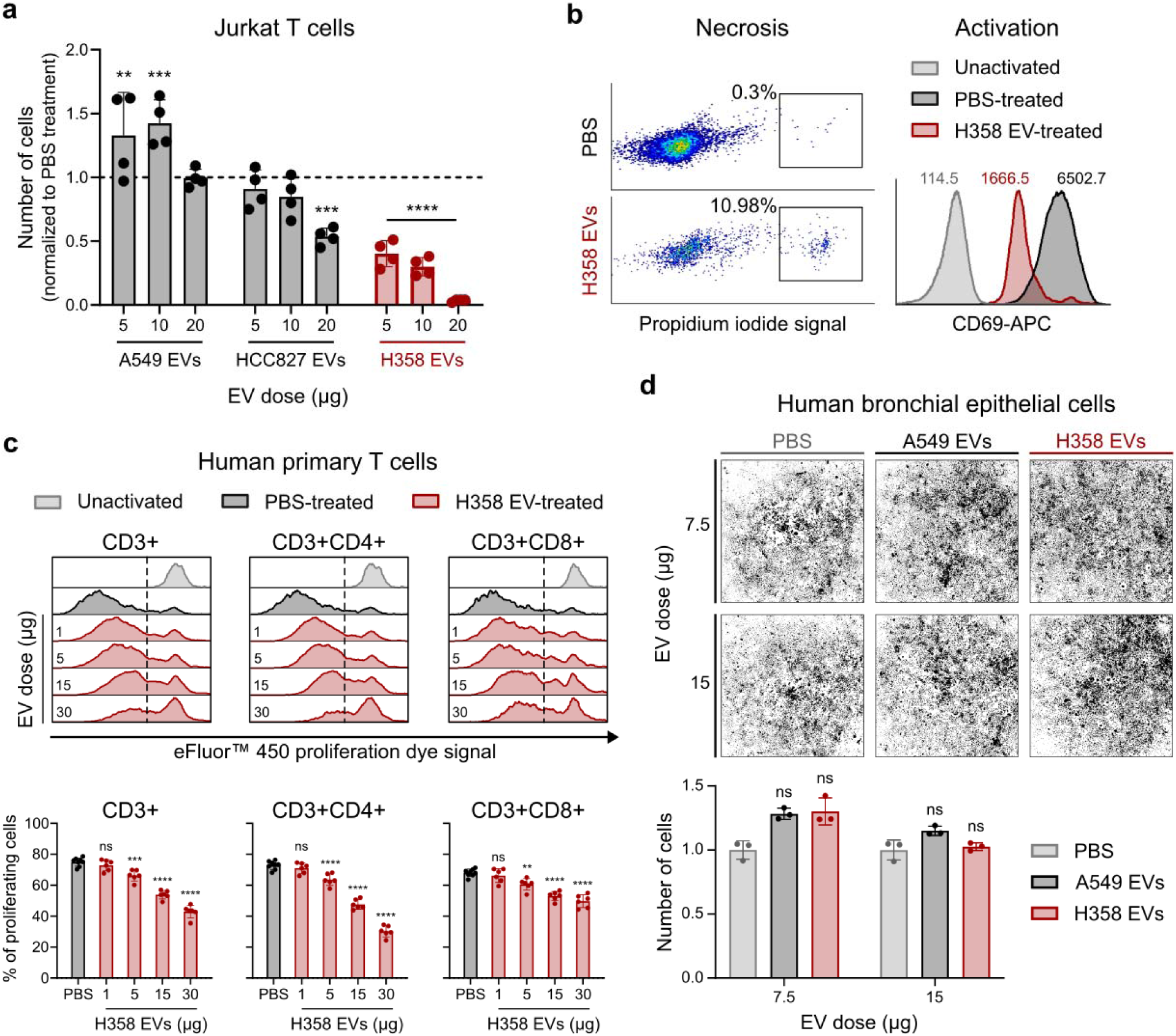
H358-derived EVs potently suppress T cell proliferation and establish a model of EV-mediated immune suppression. **a.** Jurkat T cell proliferation following treatment with EVs derived from NSCLC cell lines (A549, HCC827, and H358) at 5, 10, and 20 µg for 96 hr. Proliferation was normalized to PBS-treated Jurkat T cells (dashed line) (mean ± SD, two-way ANOVA with Dunnett’s multiple comparisons test; n=4). **b.** Left: Representative flow cytometry dot plot showing % cells positive for propidium iodide staining (necrosis marker) in Jurkat T cells treated with 25 µg of H358 EVs or PBS. Right: Representative histogram of CD69 expression (activation marker) under the same conditions. Mean values indicated above the histogram. **c.** Top: Representative histograms of eFluor™ 450 dye dilution in primary human CD3+, CD3+CD4+, and CD3+CD8+ T cells following treatment with H358 EVs for 96 hr at indicated doses. Reduced dye intensity indicated increased proliferation. Unactivated T cells and volume-matched PBS-treated T cells served as negative and positive controls, respectively. Full gating strategy shown in Fig. S6a. Bottom: Quantification of proliferating T cells (cells left of dashed line) expressed as a percentage relative to PBS control (mean ± SD, one-way ANOVA with Dunnett’s multiple comparisons test; n=6). Primary T cells from a single donor treated with six different H358 EV preps. **d.** Representative images of human bronchial epithelial cells treated with A549 or H358 EVs (7.5 and 15 µg; 96 hr) compared with PBS control. Bottom: Quantification of cell number normalized to PBS-treated control (mean ± SD, one-way ANOVA with Dunnett’s multiple comparisons test; n=3).

The immunosuppressive activity of H358-derived EVs was dose-dependent and extended to primary human T cell populations, including CD3+, CD3+CD4+ and CD3+CD8+ subsets (**Fig. 1c**). Notably, this suppressive effect was selective for T cells, as H358 EVs failed to alter the proliferation of non-tumorigenic epithelial cells (**Fig. 1d** and **Fig. S1**). These findings suggest that the immunosuppressive activity is not a universal feature of cancer-derived EVs, but rather a property of distinct EV subpopulations.

### H358 EVs suppress T cell-mediated tumor control in vivo

To determine whether H358-derived EVs suppress T cell function within the TME *in vivo*, we employed a humanized mouse model that enables functional interrogation of the human immune response. NSG mice deficient in MHC class I and II (NSG-DKO) were engrafted with human hematopoietic stem cells (hHSCs) to generate mice with a reconstructed human immune system (huNSG-DKO) (**Fig. 2a**). Successful humanization was confirmed by the presence of human CD45+ leukocytes in peripheral blood eight weeks after engraftment (**Fig. 2b** and **Fig. S2**).

**Figure 2.**
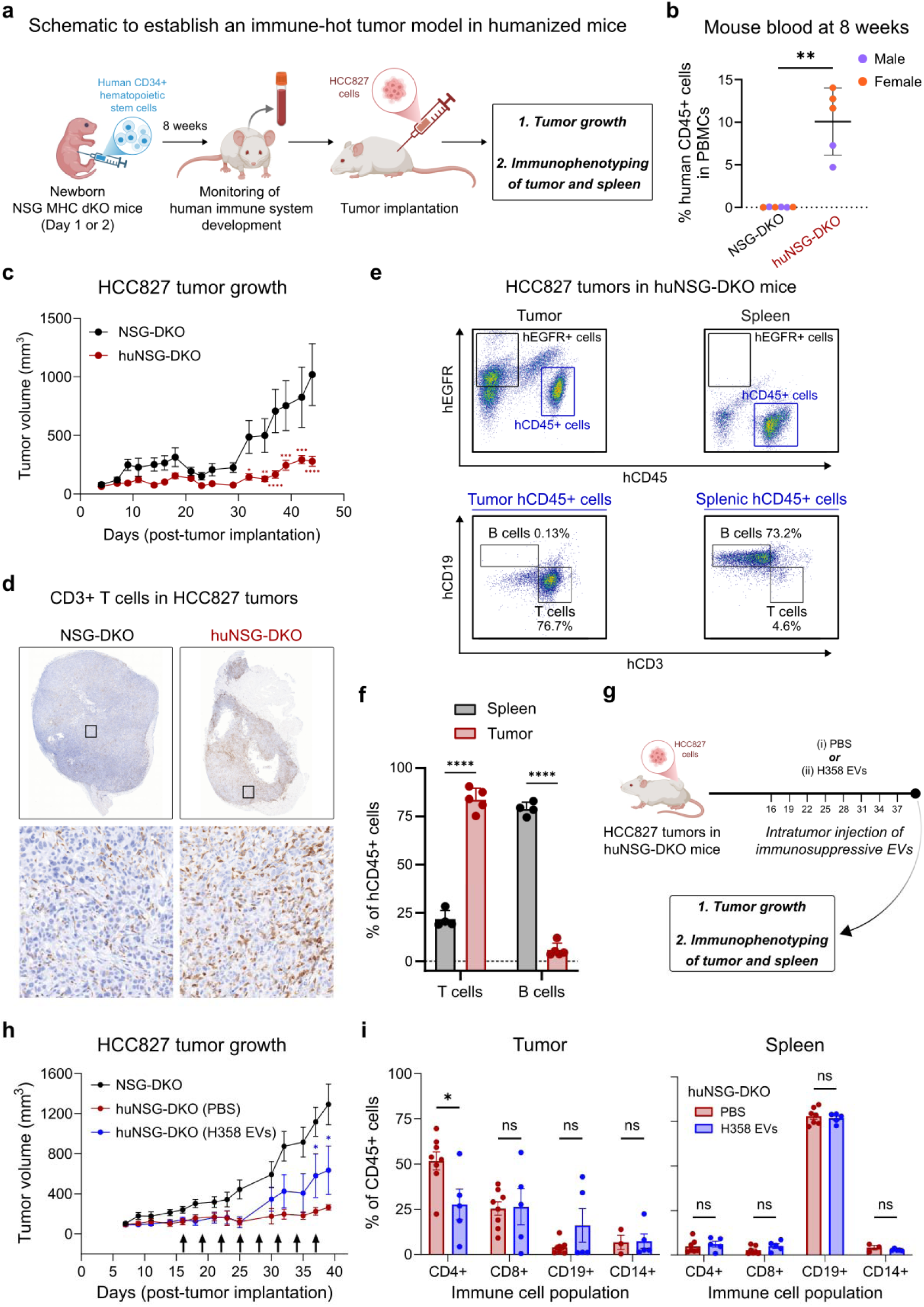
H358-derived EVs suppress T cell infiltration and promote tumor growth in a human NSCLC model. **a.** Schematic illustrating humanization of NSG-DKO mice via intrahepatic engraftment of human hematopoietic stem cells (hHSCs), followed by subcutaneous implantation of HCC827 lung cancer cells to establish an inflamed NSCLC tumor model. **b.** Frequency of human CD45+ leukocytes in peripheral blood of humanized (huNSG-DKO) mice 8 weeks after hHSC engraftment. Non-humanized NSG-DKO mice served as a control (mean ± SD; Welch’s t-test; n=6 NSG-DKO, n=5 huNSG-DKO). Full gating strategy is shown in Fig. S6b. **c.** HCC827 tumor growth in NSG-DKO and huNSG-DKO mice, measured by tumor volume (mean ± SEM; unpaired t-test; n=5 per group). **d.** Representative CD3 immunohistochemical staining of HCC827 tumors from NSG-DKO and huNSG-DKO mice. **e.** Top: Representative dot plots showing human EGFR+ tumor cells and human CD45+ immune cells in tumors and spleens of huNSG-DKO mice. Bottom: Representative dot plots depicting CD3+ T cells and CD19+ B cells within tumor-infiltrating and splenic human CD45+ populations. **f.** Quantification of T cells and B cells among tumor-infiltrating and splenic human CD45+ cells in huNSG-DKO mice (mean ± SD; unpaired t-test; n=5 for tumor, n=4 for spleen). **g.** Schematic of intratumor administration of H358-derived EVs into established HCC827 tumors in huNSG-DKO mice. **h.** HCC827 tumor growth following intratumoral injection of H358 EVs. Groups include NSG-DKO, PBS-treated huNSG-DKO, and H358 EV-treated huNSG-DKO mice (mean ± SEM; two-way ANOVA with Dunnett’s multiple comparisons test; n=10 NSG-DKO, n=8 PBS-treated huNSG-DKO, n=5 EV-treated huNSG-DKO). **i.** Flow cytometric quantification of CD4+ and CD8+ T cells, CD19+ B cells, and CD14+ monocytes within tumor-infiltrating and splenic human CD45+ cells following PBS or H358 EV administration (mean ± SD; unpaired t-test; n=5 per group). Full gating strategy is shown in Fig. S7.

We next established an immune-sensitive NSCLC tumor model by implanting HCC827 cells into NSG-DKO and huNSG-DKO mice. Tumor growth was significantly restrained in huNSG-DKO mice compared to NSG-DKO controls, reflecting efficient human immune cell-mediated tumor control (**Fig. 2c**). Consistent with this, HCC827 tumors in huNSG-DKO mice exhibited robust infiltration of human T cells (**Fig. 2d-f**), validating this system as a suitable platform to assess EV-mediated modulation of tumor-infiltrated T cells *in vivo*.

To directly test the T cell-suppressive activity of tumor-derived EVs within the TME, HCC827 tumors in huNSG-DKO mice were treated intratumorally with H358 EVs or PBS every three days (**Fig. 2g**). Administration of H358 EVs markedly accelerated tumor growth compared to PBS-treated controls (**Fig. 2h**), indicating a functional impairment of anti-tumor immune responses. Immunophenotyping of tumors and spleen at study endpoint revealed that intratumor delivery of H358 EVs significantly reduced CD4+ T cell infiltration within tumors (**Fig. 2i, left panel**). Importantly, no changes in immune cell abundance or T cell activation were observed in the spleen, demonstrating that the immunosuppressive effects of H358 EVs were localized to the tumor siter rather than systemic (**Fig. 2i, right panel**).

Collectively, these findings established that H358-derived EVs actively suppress T cell-mediated tumor control *in vivo* and validate this NSCLC model as a robust system for dissecting mechanisms of EV-driven immunosuppression within the TME.

### TGOLN2+ EVs represent a distinct, Golgi-derived EV subpopulation that mediates T cell suppression

To define the EV population responsible for T cell suppression, we first examined whether canonical tetraspanin-positive EVs mediate this effect. Although H358-derived EVs abundantly expressed CD81 and to a lesser extent CD63 (**Fig. 3a**), immunodepletion of either CD81+ or CD63+ EVs from the total EV pool did not diminish the remaining EVs from suppressing T cell proliferation (**Fig. 3b-c**). These findings indicate that the T cell-suppressive activity resides within a distinct EV subpopulation that lacks conventional tetraspanin markers.

**Figure 3.**
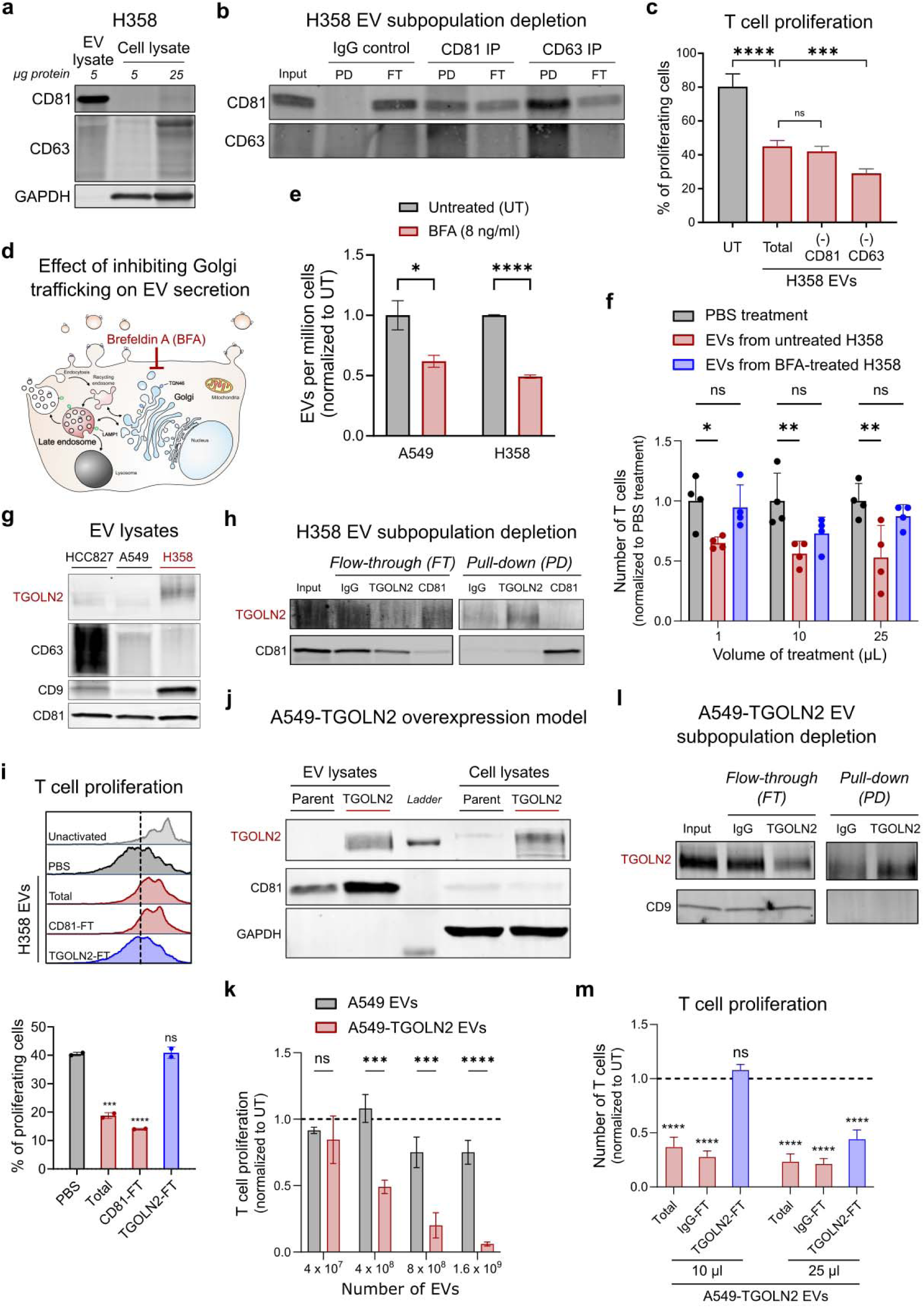
TGOLN2+, tetraspanin-low, EV subpopulation mediates T cell suppression. **a.** Representative immunoblot showing canonical EV markers CD81 and CD63 in H358 EV lysate (5 µg total protein) compared to H358 whole-cell lysate (5 and 25 µg). GAPDH confirms minimal cellular contamination of isolated EVs. **b.** Immunoblot of pull-down (PD) and flow-through (FT) fractions following CD81- and CD63-immunoprecipitation of H358 EVs. IgG immunoprecipitation served as a control. **c.** Jurkat T cell proliferation following treatment with total H358 EVs or EVs depleted of CD81+ or CD63+ vesicles (96 hr). EV input was normalized to 5 µg total protein. Proliferation was quantified by eFluor™ 450 dye dilution (mean ± SD; unpaired t-test; n=3). **d.** Schematic illustrating Brefeldin A (BFA) treatment to inhibit Golgi trafficking and assess its effect on EV secretion. **e.** Quantification of EV secretion from A549 and H358 cells following 48 hr treatment with BFA (8 ng/mL). EV number is shown per 10^6^ cells. Untreated cells served as a control (mean ± SD; unpaired t-test; n=3). **f.** Jurkat T cell proliferation following treatment with equal volumes of EVs derived from untreated or BFA-treated H358 cells. Proliferation was normalized to PBS-treated Jurkat T cells (mean ± SD; unpaired t-test; n=4). **g.** Immunoblot showing expression of the trans-Golgi marker TGOLN2 and canonical EV markers in EV lysates (10 µg total protein) from HCC827, A549, and H358 cells. **h.** Immunoblot of PD and FT fractions following TGOLN2 or CD81 immunoprecipitation of H358 EVs. IgG immunoprecipitation served as a control. **i.** Top: Representative histogram of primary human T cell proliferation following treatment with total H358 EVs or EVs depleted of CD81+ or CD63+ vesicles (96 hr), assessed by eFluor™ 450 dye dilution. Bottom: Quantification of proliferating T cells (cells left of dashed line) relative to PBS-treated controls (mean ± SD; one-way ANOVA with Dunnett’s post hoc, n=6). **j.** Immunoblot of TGOLN2 in EV and whole-cell lysates from TGOLN2-overexpressing A549 cells (A549-TGOLN2) and parent A549 cells. CD81 and GAPDH confirm EV enrichment and purity. **k.** Jurkat T cell proliferation following treatment with equal numbers of EVs from parental A549 or A549-TGOLN2 cells. Proliferation normalized to PBS-treated controls (mean ± SD; unpaired t-test; n=4). **l.** Immunoblot of PD and FT fractions following TGOLN2-immunoprecipitation of EVs derived from A549-TGOLN2 cells. CD9 is shown as a canonical EV marker. IgG served as a control. **m.** Jurkat T cell proliferation following treatment with total A549-TGOLN2 EVs, mock-depleted EVs (IgG), or TGOLN2-depleted EVs (96 hr; 10 and 25 µL input volumes). Proliferation normalized to PBS-treated Jurkat T cells (mean ± SD; unpaired t-test; n=4). *p<0.05, **p<0.01, ***p<0.001, ****p<0.0001

We next investigated whether this unconventional EV population originates from a non-canonical intracellular trafficking pathway. Pharmacological inhibition of Golgi trafficking using non-toxic doses of Brefeldin A (**Fig. S3a**) significantly reduced overall EV secretion from both A549 and H358 cells (**Fig. 3d-e**). Importantly, EVs isolated from Brefeldin A-treated H358 cells were deficient in their ability to suppress T cell proliferation when normalized for EV input (**Fig. 3f**), implicating a Golgi-dependent EV biogenesis pathway in T cell suppression.

Consistent with this hypothesis, immunoblotting of Golgi-associated proteins revealed that H358 EVs were highly enriched for TGOLN2, a trans-Golgi network (TGN) marker, compared to EVs derived from immune-neutral NSCLC models (**Fig. 3g and Fig. S3b**). Despite co-expression of CD81 and CD63 within the total EV population (**Fig. 3b**), TGOLN2 expression exhibited minimal overlap with CD81 in co-immunoprecipitation experiments (**Fig. 3h**), further supporting the existence of a discrete TGOLN2+ EV subpopulation. This population was validated as functionally relevant as depletion of the TGOLN2+ EVs from H358-derived EVs significantly abrogated the T cell-suppressive activity of the EVs (**Fig. 3i**), identifying TGOLN2+ EVs as the primary mediators of T cell suppression.

To determine whether TGOLN2 was merely a marker of the T cell suppressive EVs or functionally contributes to T cell suppressive EV biology, we ectopically expressed TGOLN2 in the immune-neutral A549 NSCLC model. TGOLN2 was abundant in EVs isolated from A549-TGOLN2 overexpressing cells (**Fig. 3j**) and, notably, conferred potent T cell suppressive activity to A549-derived EVs (**Fig. 3k**). As observed in the H358 model, TGOLN2+ EVs isolated from A549-TGOLN2 overexpressing cells exhibited minimal overlap with the canonical tetraspanin CD9 (**Fig. 3l**). Depletion of the TGOLN2+ subpopulation of EVs rescued T cell proliferation (**Fig. 3m**), confirming that TGOLN2 expression is sufficient to drive the emergence of immunosuppressive EVs.

Collectively, these data identify a previously unrecognized TGOLN2+ EV subpopulation that originates from the Golgi, lacks canonical tetraspanin markers, and potently suppresses T cell proliferation and function. This work reveals a non-classical EV biogenesis pathway as a critical regulator of tumor-induced immunosuppression.

### TGOLN2 expression correlates with poor prognosis and an immunosuppressive TME

Our findings indicate that TGOLN2 overexpression in NSCLC promotes tumorigenesis by enhancing the capacity of cancer cells to suppress T cell activity, thereby facilitating immune evasion. To evaluate the clinical relevance of this mechanism, we performed bioinformatic analysis of patient data from The Cancer Genome Atlas (TCGA), examining relationships between TGOLN2 copy number, transcript abundance, and protein expression with multiple survival endpoints. In a pan-cancer analysis, patients with elevated TGOLN2 copy number exhibited significantly worse overall survival, disease-specific survival, disease-free interval, and progression-free interval compared to patients with lower copy number (**Fig. 4a**). Among these metrics, the association with progression-free interval was the most pronounced. Consistent with these findings, higher *TGOLN2* transcript levels were also significantly associated with poorer progression-free interval across multiple cancer types, including lung adenocarcinoma (LUAD), lung squamous cell carcinoma (LUSC), glioblastoma, lower-grade glioma, and bladder cancer (**Fig. 4b**). This inverse relationship between TGOLN2 expression and patient outcome was further corroborated at the protein level using data from the Pediatric Brain Tumor Atlas (**Fig. 4c**). Collectively, these analyses demonstrate that elevated TGOLN2 expression is broadly associated with poor prognosis across diverse malignancies. Given our experimental evidence that TGOLN2 promotes T cell suppression in NSCLC, we next asked whether TGOLN2 expression correlates with features of an immunosuppressive TME in patient tumors. In both LUAD and LUSC, TGOLN2 expression negatively correlated with infiltration of CD4+ Th1 and CD8+ naïve T cells (**Fig. 4d**). Extending this analysis across TCGA cancer types revealed that in at least 20 out of the 32 malignancies, high TGOLN2 expression was associated with a predominantly immunosuppressive TME signature. This signature was characterized by reduced abundance of CD8+ T cells, activated NK cells, follicular helper T cells, and regulatory T cells, together with increased infiltration of M2 macrophages (**Fig. 4e**). *p<0.05, **p<0.01, ***p<0.001

**Figure 4.**
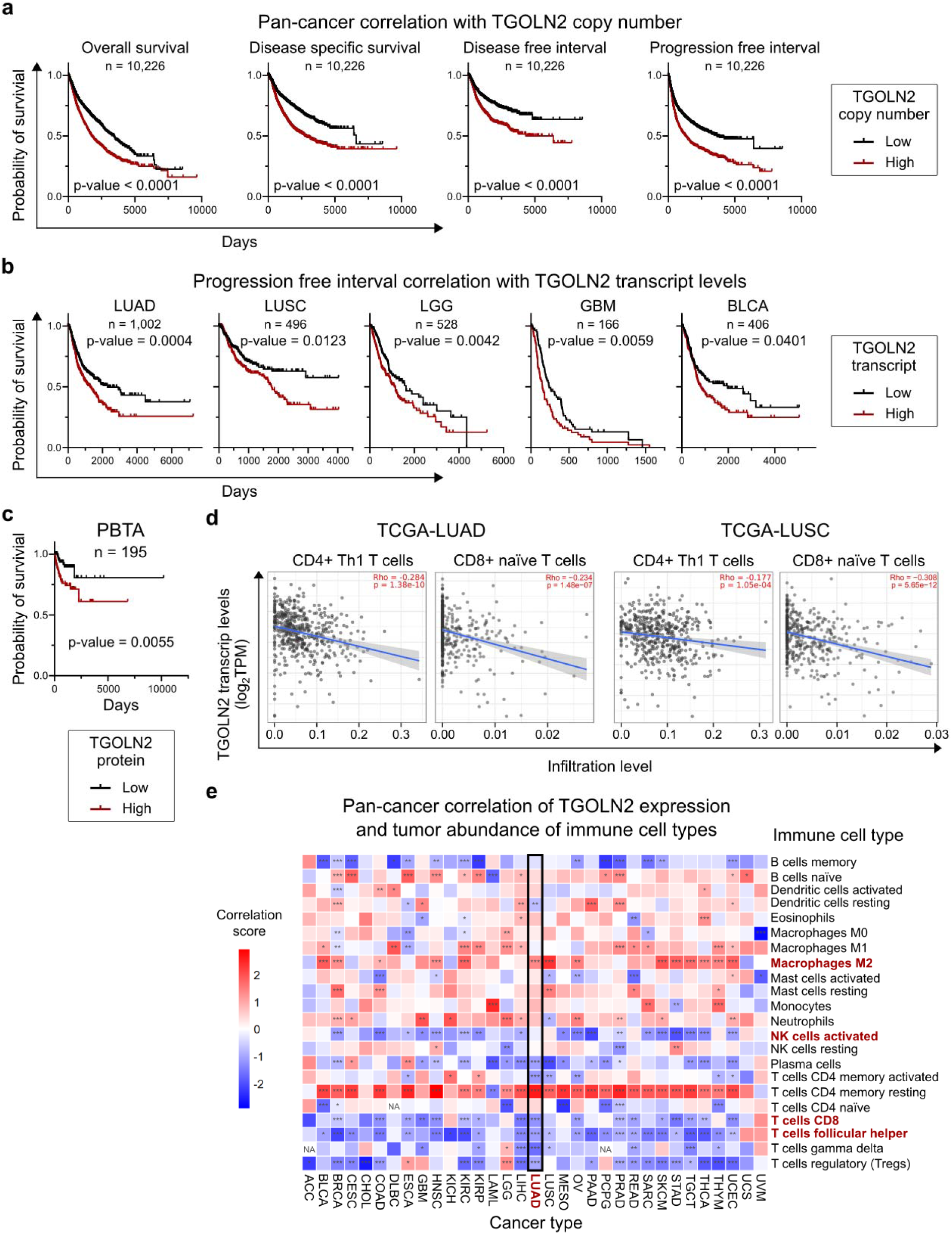
TGOLN2 overexpression associates with poor clinical outcomes and an immunosuppressive TME across cancers. **a.** Pan-cancer analysis of TGOLN2 copy number and its association with overall survival (OS), disease-specific survival (DSS), disease-free interval (DFI), and progression-free interval (PFI) in TGCA cohorts. **b.** Association between TGOLN2 transcript expression and PFI in LUAD (lung adenocarcinoma), LUSC (lung squamous carcinoma), LGG (lower grade glioma), GBM (glioblastoma), and BLCA (bladder cancer) patient datasets. **c.** Correlation of TGOLN2 protein expression with overall survival in the Pediatric Brain Tumor Atlas (PBTA). **d.** Correlation betweenTGOLN2 transcript levels and inferred tumor infiltration of CD4+ Th1 cells and CD8+ naïve T cells in LUAD and LUSC cohorts. **e.** Pan-cancer correlation of TGOLN2 transcript levels with inferred immune cell infiltration across tumor types. Red indicates positive correlation and blue indicates negative correlation (Pearson correlation; n = number of tumor samples per cancer type). *p<0.05, **p<0.01, ***p<0.001

### TGOLN2+ EVs arise through a multivesicular body-like pathway linked to the Golgi and exhibit distinct immunomodulatory properties

Because TGOLN2+ EVs had minimal overlap with conventional tetraspanin markers, we next sought to define the biogenesis of TGOLN2+ EVs and cargo within this new EV subpopulation. To visualize TGOLN2+ EV biogenesis, we generated an electron microscopy-based genetic model in which TGOLN2 was fused to APEX2 (**Fig. S4**), an engineered peroxidase widely used for ultrastructural labeling (**Fig. 5a-b** and **Fig. S5a**). As expected, APEX2 fused to a mitochondrial matrix protein (mito-APEX2) selectively labeled mitochondria. In contrast, TGOLN2-APEX2 (TAPEX) localized to the membrane of intraluminal vesicles within multivesicular body (MVB)-like structure (**Fig. 5c**). Because canonical MVBs typically originate from late endosomes, we next examined whether TAPEX-labeled structures colocalized with LAMP1, a marker of late endosomes. Notably, minimal colocalization was observed between TAPEX and LAMP1 (**Fig. S5b**), suggesting that TGOLN2+ MVB-like structures are unlikely to arise from the conventional late endosomal pathway and instead may originate from a Golgi-associated compartment.

**Figure 5.**
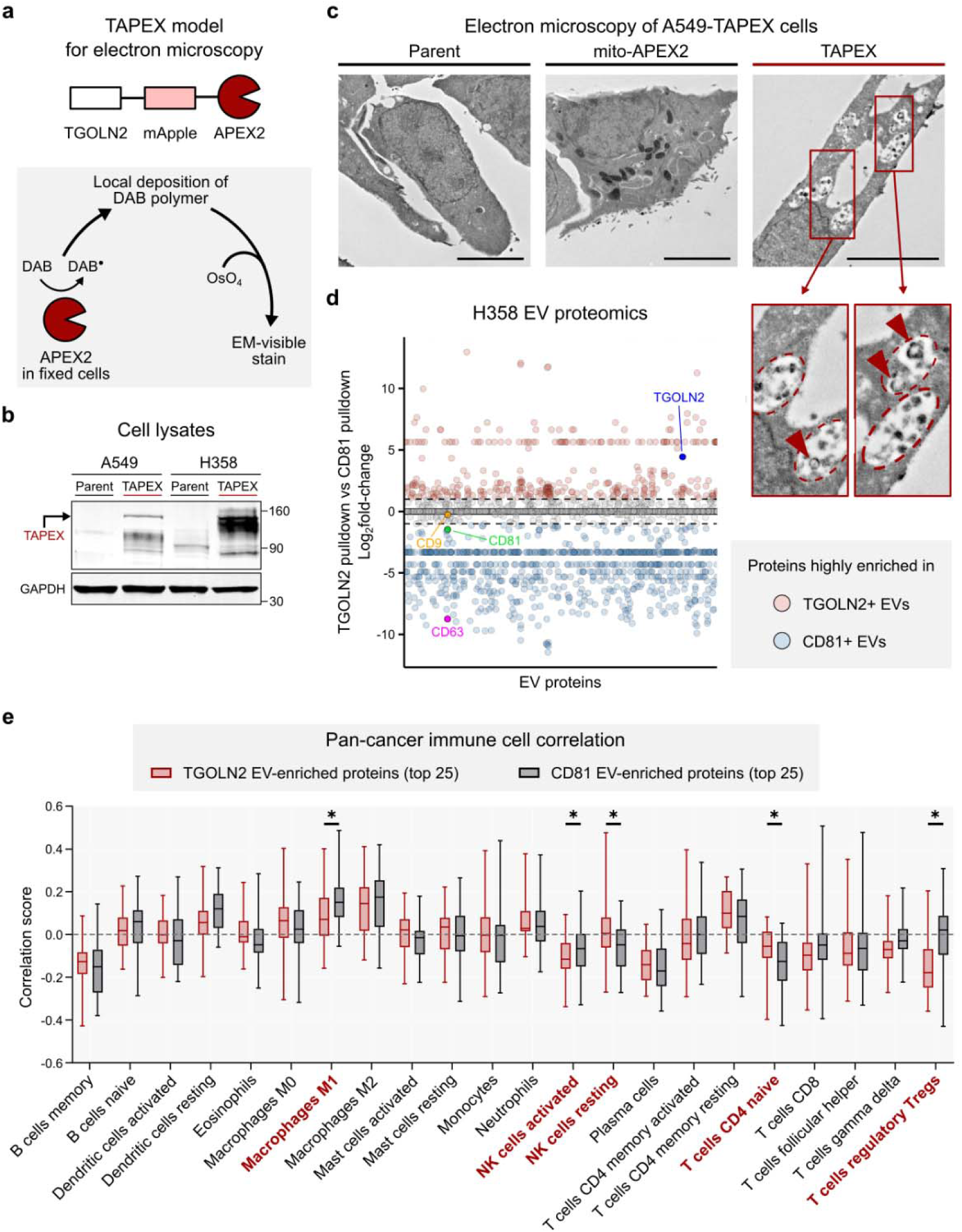
TGOLN2+ EVs arise from multivesicular body formation and exhibit a proteomic signature associated with immunosuppressive TMEs. **a.** Schematic of APEX2-tagged TGOLN2 (TAPEX) construct used for ultrastructural localization by electron microscopy. **b.** Immunoblot confirming TAPEX expression in A549-TAPEX and H358-TAPEX cells compared with parental controls. **c.** Representative electron micrographs demonstrating electron-dense APEX2 signal in TAPEX-expressing A549 cells. Parent A549 cells and A549 cells expressing mito-APEX2 (mitochondrial matrix-localized APEX2) served as negative and positive controls, respectively. **d.** Comparative proteomic analysis of TGOLN2+ and CD81+ EVs isolated from H358 EVs via immunoprecipitation. Protein enrichment is shown as log_2_ fold-change (FC) (TGOLN2+ EVs vs. CD81+ EVs). Red indicates proteins enriched in TGOLN2+ EVs (log_2_ FC > 1), and blue indicates enrichment in CD81+ EVs (log_2_ FC < −1). **e.** Correlation of tumor immune infiltration with the mean expression of top 25 proteins enriched in TGOLN2+ EVs or CD81+ EVs across 33 cancer types. Positive correlation indicates higher expression associated with increased immune infiltration. Negative correlation indicates higher expression associated with decreased immune infiltration (mean ± SEM; paired t-test; n=33 cancer types). *p<0.05, **p<0.01, ***p<0.001, ****p<0.0001

We next investigated the molecular cargo of TGOLN2+ EVs, by identifying proteins selectively enriched in TGOLN2+ EVs compared to CD81+ EVs (**Fig. 5d**). Ontology analysis of the top 100 proteins enriched in TGOLN2+ EVs revealed significant enrichment for the molecular function term “immunoglobulin binding” and the cellular component “spliceosome” (**Supplementary Bioinformatics**, Sheet B). In contrast, proteins enriched in CD81+ EVs were significantly associated with “cell adhesion” and “G protein activity” molecular functions and the “cell surface” cellular component (**Supplementary Bioinformatics**, Sheet C). Both EV populations had significant enrichment for the “extracellular exosome” cellular component, confirming their identity as EVs. Given the distinct proteomic composition of TGOLN2+ EVs relative to CD81+EVs, we next asked whether these EV subpopulations were correlated with differential immunomodulatory effects within the TME. Immune cell correlation analysis revealed that TGOLN2+ EVs were more strongly associated with regulatory T cells, whereas both TGOLN2+ and CD81+ EVs had comparable associations with other immune populations, including CD8+ T cells, activated NK cells, follicular helper T cells, and M2 macrophages (**Fig. 5e**).

Together, these findings demonstrate that TGOLN2⁺ EVs represent a biologically distinct EV subpopulation that arises through a non-canonical, Golgi-associated MVB-like pathway and carries a unique protein cargo linked to immunomodulatory functions. These results underscore how EV heterogeneity enables cancer cells to shape the tumor microenvironment through both shared and specialized mechanisms.

## DISCUSSION

Tumor-derived EVs have emerged as important mediators that contribute to resistance to cancer immunotherapy^9^. However, significant EV heterogeneity, poor understanding of EV biogenesis, and limited models to study EV biology have hindered the identification of specific EV subpopulations responsible for immunosuppression and immunotherapy resistance. In this study, we developed human-relevant NSCLC models to investigate EV-mediated immunosuppression both *in vitro* and *in vivo*. Using these models, we identified a distinct subpopulation of cancer-derived EVs that potently suppress T cell activation and proliferation. Our findings identify TGOLN2 – previously characterized primarily as a TGN marker – as a potential oncogene and implicate its overexpression as a key mechanism driving EV-mediated T cell suppression, thereby promoting an immunosuppressive TME and potentially contributing to immunotherapy resistance.

We first performed an *in vitro* screen and identified H358 as a human NSCLC model that exhibits strong EV-mediated T cell suppression (**Fig. 1**). To study EV-mediated immunosuppression in a human-relevant context, we established a humanized NSG-DKO (huNSG-DKO) mouse model lacking mouse major histocompatibility complex (MHC)-I and MHC-II expression. This model supports robust development of the human immune system following engraftment with human hematopoietic stem cells (hHSCs) (**Fig. 2**). Although immune reconstruction in this model is less extensive than in some advanced humanized mouse systems, the absence of mouse MHC-I/II allows human HLA-restricted immune system development, likely through extrathymic education, enabling the study of physiologically relevant human immunobiology. Using this platform, we established HCC827 as an immune-sensitive human NSCLC tumor model that exhibited reduced tumor growth and inflamed TME characterized by high T cell infiltration. These findings were consistent across multiple independent studies using huNSG-DKO mice engrafted with hHSCs from different donors (**Fig. 2**). Together, these models provide a robust framework for studying EV-mediated modulation of the TME in NSCLC and may be adapted to other cancer types.

Using these complementary models, we evaluated the impact of H358-derived EVs on the inflamed TME of HCC827 tumors. H358 EVs reduced intratumoral CD4+ T cell infiltration (**Fig. 2**). Because secreted EVs can enter systemic circulation and affect distant tissues, we also assessed T cell abundance in the spleen of the same mice. We observed no significant effects on splenic T cells, suggesting that the immunosuppressive activity of H358 EVs is tumor localized. Nonetheless, potential effects on circulating T cells or immune cells in adjacent tissues cannot be excluded and warrant further investigation. Additionally, EVs are known to modulate multiple cell types. Consistent with this, we previously demonstrated that H358 EVs H358 EVs can increase invasive capacity of non-tumorigenic epithelial cells^14^. Thus, H358 EVs may influence additional immune and non-immune populations within the TME, underscoring the need for comprehensive single-cell and spatial omics approaches to define their broader biological impact.

EV heterogeneity remains a major challenge in EV biology, largely due to lack of definitive markers that distinguish functionally distinct EV subpopulations. To address this limitation, we used the H358 model to perform immunoprecipitation-based depletion of specific EV subpopulations and subsequently assessed the immunomodulatory activity of the remaining EVs (**Fig. 3**). This strategy enabled functional attribution of activity to defined EV fractions and identified TGOLN2+ EVs as the specific subpopulation responsible for T cell suppression.

Notably, TGOLN2+ EVs exhibited minimal overlap with the canonical EV markers, including CD81, CD63, and CD9, suggesting that certain functionally-relevant EV subpopulations may lack these commonly used markers. These findings highlight an important limitation of relying exclusively on traditional tetraspanins markers to define biologically active EV populations.

Leveraging recent advances in genetically encoded electron microscopy (EM) tags^14,15^, we next developed a genetic EM-based model to visualize TGOLN2+ EVs biogenesis. These studies revealed that TGOLN2+ EVs originate from the TGN through the formation of multivesicular body-like structures. EVs have historically been described as originating either from plasma membrane budding or from late endosomal multivesicular bodies^10^, with more recent evidence supporting mitochondrial-derived pathways^11^. However, the contribution of the Golgi apparatus to EV biogenesis has remained largely unexplored. Our findings demonstrate, for the first time, that the Golgi compartment plays an active role in EV biogenesis and contributes directly to EV-mediated immunosuppressive biology in NSCLC.

Although isolation and functional testing of exclusively TGOLN2⁺ EVs was not technically feasible due to limitations in capturing and releasing intact immunoprecipitated EVs, the combined evidence from selective depletion experiments and TGOLN2 overexpression studies strongly supports TGOLN2⁺ EVs as immunomodulatory. Future studies should prioritize direct functional evaluation of the TGOLN2⁺-only subpopulation as enabling technologies continue to advance.

Lastly, we identified TGOLN2 overexpression as the underlying mechanism driving increased release of TGOLN2+ EVs and enhanced T cell suppression. TGOLN2 is a type I integral membrane protein that localizes to the TGN at steady state^16,17^, but rapidly cycles between the TGN and the plasma membrane^18^. TGOLN2 is known to mediate sorting of soluble cargo into CARTS (<u>car</u>riers of the <u>T</u>GN to the cell <u>s</u>urface). Consistent with this role, a recent study demonstrated that TGONL2 regulates the constitutive secretion of PAUF^19^, a soluble protein highly expressed in pancreatic ductal adenocarcinoma^20^ that promotes immune evasion by enhancing recruitment and function of myeloid-derived suppressor cells (MDSCs)^21^. Together with our findings, these data suggest that TGOLN2 overexpression may simultaneously promote secretion of T cell-suppressive EVs and MDSC-activating soluble factors, thereby reinforcing an immunosuppressive TME. Supporting this model, analysis of TCGA patient datasets revealed that elevated TGOLN2 expression correlates with immunosuppressive TME signatures across 20 cancer types with poor prognosis in at least six cancer types (**Fig. 4**). Notably, in prostate adenocarcinoma and kidney clear-cell carcinoma, higher TGOLN2 expression was associated with immunosuppressive TME signatures yet correlated with improved prognosis, suggesting cancer type-specific roles for TGOLN2 that warrant further investigation.

In summary, we developed physiologically relevant *in vitro* and *in vivo* models to study EV-mediated immunosuppression in NSCLC. Using these systems, we dissected EV heterogeneity and identified TGOLN2+ EVs as a distinct cancer-derived EV subpopulation that originates from the TGN and potently suppresses T cell function. We further identified TGOLN2 overexpression as the underlying mechanism and propose TGOLN2 as a candidate oncogene that promotes an immunosuppressive TME in NSCLC and potentially other cancers. These findings provide new insights into the biology of immunosuppressive TMEs and reveal therapeutic opportunities to target EV-mediated immune evasion and immunotherapy-resistant NSCLC.

## METHODS

### Cell lines

A549 (CCL-185™, RRID:CVCL_0023), HCC827 (CRL-2868™, RRID:CVCL_2063) and H358 (CRL-5807™, RRID:CVCL_1559) were purchased from ATCC, and Jurkat T cells (RRID:CVCL_0065) were provided by Dr. Kazemian (Purdue University). All cell lines were cultured in complete RPMI [RPMI 1640 (SH30027FS, Fisher Scientific Inc.) supplemented with 10% FBS (fetal bovine serum, S11150, Bio-Techne) and 1X penicillin/streptomycin (SV30010, Fisher Scientific Inc.)]. Cell lines were monitored monthly for lack of mycoplasma using the MycoAlert Mycoplasma Detection Kit (LT07-418, Lonza). For primary T cell experiments, human peripheral blood mononuclear cells (PBMCs) were used (see protocol below for generating PBMCs). Human PBMCs were cultured in the same media as mentioned above, except where otherwise noted.

### Extracellular vesicle isolation and characterization

Extracellular vesicles were isolated from cell culture supernatant as described before^22^. Briefly, when cells reached ∼80% confluency, normal culture media was replaced with media made with EV-depleted FBS. After 48 hours, media was collected and centrifuged at 2,000xg followed by centrifugation at 10,000xg to remove cells, cell debris, and large apoptotic EVs. The media was then filtered through a 0.2 µm filter to remove large particles and inadvertent contaminants and centrifuged at 110,000xg to pellet EVs that were enriched for exosomes. The resulting EV pellet was washed in 1X PBS (SH30256FS, Fisher Scientific Inc.) and centrifuged again at 110,000xg. The final pellet was resuspended in PBS and stored at −80 °C until further use. The isolated EVs were extensively characterized as per the guidelines of the International Society for Extracellular Vesicles using transmission electron microscopy, nanoparticle tracking analysis, and western blotting for EV-enriched tetraspanins (CD9, CD81, and CD63) and cell-enriched proteins (GP96)^23^.

For transmission electron microscopy, samples were deposited onto glow-discharged carbon-coated 200-mesh copper grids (Electron Microscopy Sciences, CF200-Cu) in the presence of 2% PTA (phosphotungstic acid) at a 1:1 ratio. After 30 s, excessive moisture was wicked off using filter paper. The grid was air-dried and imaged using Tecnai T12 transmission electron microscope at 100 keV using a Gatan US1000 2 KL×L2 K CCD camera.

### Protein extraction and western blotting

Cells were lysed using RIPA buffer [Tris-HCl (pH 8.0, 50 mM), NP-40 (1%), Sodium chloride (150 mM), Sodium deoxycholate (0.5%), SDS (0.1%), ddH2O (up to 100 mL)] in the presence of 1X protease inhibitor (PIA32955, Thermo Fisher Scientific Inc.) and 1X phosphatase inhibitor cocktail (PIA32957, Thermo Fisher Scientific Inc.). EVs were lysed using Laemmli’s SDS-Sample buffer (NC9409160, Fisher Scientific Inc.). Protein concentration was measured using the Pierce™ BCA Protein Assay kit (23225, Thermo Fisher Scientific Inc.). Cell and EV lysate were resolved on 4-20% TGX™ gels (5671093, Bio-Rad Laboratories Inc.) and transferred to Immobilon™-FL PVDF (polyvinylidene difluoride) membrane (IPFL00005, Millipore). The PVDF membrane was incubated in Intercept™ blocking buffer (927-70001, LI-COR Biotech LLC) for 1 hr at room temperature. After blocking, the membrane was incubated overnight in the indicated primary antibody at 4 °C followed by corresponding secondary antibody incubation for 1 hr at room temperature. The membrane was washed and scanned using Odyssey CLx imaging system and Image Studio™ software (LI-COR Biotech, LLC) with scan resolution set to “169 µm”, scan quality set to “high” or “highest”, and focus offset set to “0.0 mm”. Antibodies used for western blotting and other experiments are indicated in **Table S1**.

### Confocal microscopy

A day before seeding cells, glass coverslips were inserted in a 24-well plate and coated with 8 µg/cm^2^ type IV collagen (C5138-100MG, Sigma-Aldrich Inc.). Equal numbers of cells were seeded on collagen-coated coverslips followed by culturing for 48 hr with or without any treatment as indicated. After 48 hr, cells were prepared for immunostaining. All washes and staining steps were performed at room temperature with gentle rocking. Cells were washed 5L×L10 min with 200 µL PBS, incubated for 15 min with 200 µL 4% paraformaldehyde (Fisher Scientific Inc., cat. no. AAJ61899AK) to fix, washed 2L×L10 min with PBS, incubated with 100 µL of 0.5% Saponin (55-825-5, Fisher Scientific Inc.) in PBS for 10 min to permeabilize, washed 4L×L5 min with PBS, and incubated in blocking buffer (1% BSAL+L0.1% Saponin in PBS) for 30 min. Cells were then incubated with primary antibody diluted in 100 µL of blocking buffer for 3 h, washed 5L×L10 min with 200 µL blocking buffer, incubated with fluorophore-conjugated secondary antibody at 4 µg/mL and Hoechst 33342 (62249, Thermo Fisher Scientific Inc.) at 10 µg/mL in blocking buffer for 2 h and washed 5L×L10 min with PBS. Coverslips were then mounted on glass slides in Prolong™ Glass Antifade reagent (P36982, Fisher Scientific Inc.). Confocal fluorescence images were acquired using a Nikon A1R-MP multiphoton confocal microscope with a 40X objective (Nikon Inc.), using standard excitation and emission fluorescence channels for AlexaFluor 488 and Hoechst. The images were acquired and analyzed using the Nikon NIS Elements software in the .nd2 format. The acquisition settings were 2 K x 2 K resolution (pixels) with a scanning frame rate of 1/16 s. All images were set to the same display lookup table (LUT) settings before exporting the files and mean channel intensities.

### TAPEX validation and electron microscopy

APEX2 activity was validated by light microscopy, as described here^15^. Briefly, A549 parent and A549-TAPEX stable clone cells were seeded at 50,000 cells per well in a 12-well plate. For mito-V5-APEX2 (positive control), A549 parent cells were transfected with the plasmid vector (Addgene, 72480, RRID:Addgene_72480). The media was changed for all cells after 6 hr of transfection. After 48 hr of transfection, cells were fixed in warm (30-37 °C) 2.5% glutaraldehyde solution. The solution was immediately removed and replaced with fresh 2.5% glutaraldehyde solution and incubated at room temperature for 5 min. The cells were transferred on ice for 60 min and then washed three times for 1 min each in cold 1X PBS. After PBS washes, cells were incubated with cold 20 mM glycine solution for 5 min on ice. A fresh solution containing diaminobenzidine (DAB) at 0.5 mg/mL and 10 mM H_2_O_2_ was prepared (1X DAB solution). Cells were incubated with 1X DAB solution on ice until light brown staining was visible under a light microscope. Once brown staining was visible, 1X DAB solution was removed and cells were washed 3 x 1 min with 1X PBS.

For electron microscopy, A549 parent, A549-TAPEX, and A549 parent cells transfected with mito-V5-APEX2 were processed as described above with two modifications^15^. Cells were incubated with 1X DAB solution on ice for 2 hr and 1X sodium cacodylate solution (100 mM sodium cacodylate, pH 7.4, with 2 mM calcium chloride) was used instead of 1X PBS. Following DAB incubation and washes, cells were incubated for 30 min with a fresh solution of 2% (wt/vol) OsO_4_ in cold 1X sodium cacodylate. The OsO4 solution is gently removed and transferred to a waste container containing sodium sulfite quenching solution (0.5M sodium sulfite in cold 1X sodium cacodylate). Cells were washed 5 x 2 min with ice-cold water followed by incubating the cells in cold 2% (wt/vol) urany acetate solution at 4 °C overnight in the dark. Cells were washed 5 x 2 min with ice-cold water. Cells were brought to room temperature, washed in distilled water, and then carefully scraped off the plastic, resuspended, and centrifuged at 700g for 1 min to generate a cell pellet and submitted to Purdue Electron Microscopy Center for dehydration and resin embedding process. The supernatant was removed, and the pellet was dehydrated in a graded ethanol series (50%, 75%, 90%, 95%, 100%, 100%, 100%), for 10 min each time, then infiltrated in EMBED-812 (Electron Microscopy Sciences) using 1:1 (v/v) resin and anhydrous ethanol overnight, followed by two changes into 100% resin before letting sit overnight. Finally, the sample was exchanged once more with 100% resin before transfer to fresh resin and polymerization at 60 °C for 48 h. Embedded cell pellets were cut with a diamond knife into 50-nm sections and imaged on a FEI-Tecnai T12 transmission electron microscope operated at 80 kV. Images were acquired using the equipped Gatan MSC794 CCD camera at 2K x 2K resolution.

### Cell proliferation assay

Human bronchial epithelial cells (HBECs) were seeded at 5,000 cells per well in a 96-well plate. After 24 hr, cells were treated with different doses of EVs as indicated. After 96 hr of EV treatment, cells were fixed and stained using Differential Quik^®^ III staining kit (26419-16; Polysciences, Inc.) as per manufacturer instructions. The stained cells were imaged using Olympus IX73 microscope at 4X magnification. The imaging was performed over the entire well with three wells per sample. The number of cells was quantified using ImageJ (RRID:SCR_003070). Images were converted to RGB stack and for the highest contrast stack, threshold was adjusted to “10-150”. The number of cells was quantified by using the “Analyze particles” function with size (pixel^2^) set to “25-Infinity” and other settings as default (**Fig. S1b**). The data was compiled and analyzed using GraphPad Prism v10 (GraphPad Software, LLC, RRID:SCR_002798).

### EV subpopulation depletion by immunoprecipitation

Before immunoprecipitation, 10 µg of indicated antibody was cross-linked to 200 µL of Dynabeads™ Protein G (10003D, Thermo Fisher Scientific Inc.) using the protocol described here^24^. EVs (equivalent to 20 µg EV protein) were incubated with Dynabeads™-crosslinked antibody overnight on a tube rotator (10136-084, VWR International, LLC) at 4 °C. Incubated EVs were precipitated by placing the tubes in a magnetic separation rack (S1506S, New England Biolabs) and the supernatant was collected as the flow-through (FT) fraction. The pellet was resuspended in 1X PBS (SH30256FS, Fisher Scientific Inc.) and was collected as the pull-down (PD) fraction. To confirm successful immunoprecipitation, western blotting was performed using equal volumes of FT and PD fractions. FT fraction (unbound EVs) was used for further downstream functional assays.

### EV subpopulation proteomics

#### Sample preparation

The pull down fractions from EV immunoprecipitation (see above) were used to conduct proteomics analysis. The prepared EV samples were submitted to Tymora Analytical Operations (West Lafayette, IN). Samples were lysed and denatured to extract proteins using phase-transfer solution lysis buffer, as described before^25^. The proteins were reduced and alkylated by incubation in 10 mM TCEP and 40 mM CAA for 10 min at 95 °C. The extracted proteins were captured onto magnetic beads using a one-pot procedure (as described previously^26^) by adding acetonitrile to 70% final concentration and incubating 10 min at room temperature. The beads were then washed three times with 70% acetonitrile and excess acetonitrile solution was allowed to dry. The proteins on beads were digested with Lys-C (Wako) at 1:100 (wt/wt) enzyme-to-protein ratio for 1 hr at 37 °C in 50 µL of 50mM triethyl ammonium bicarbonate. Trypsin was added to a final 1:50 (wt/wt) enzyme-to-protein ratio for a 3 hr digestion at 37 °C. To recover the peptides, acetonitrile was added to each sample to a final concentration of 60% to capture the enzymes onto beads. The peptide samples were recovered in the supernatant and dried completely in a vacuum centrifuge and stored at −80°C. A portion of each sample was used for peptide quantitation using Pierce Colorimetric Peptide Quantitation Kit (Thermo Fisher Scientific Inc.).

#### LC-MS/MS analysis

The dried peptide samples were dissolved to a concentration of 0.1 µg/µL in 0.05% trifluoroacetic acid with 3% (vol/vol) acetonitrile. Each sample (5 μL) was injected into an Ultimate 3000 nano UHPLC system (Thermo Fisher Scientific Inc.). Peptides were captured on a 2-cm Acclaim PepMap trap column and separated on a 50-cm column packed with ReproSil Saphir 1.8 μm C18 beads (Dr. Maisch GmbH). The mobile phase buffer consisted of 0.1% formic acid in ultrapure water (buffer A) with an eluting buffer of 0.1% formic acid in 80% (vol/vol) acetonitrile (buffer B) run with a linear 90-min gradient of 6–30% buffer B at a flow rate of 300 nL/min. The UHPLC was coupled online with an Exploris 480 mass spectrometer (Thermo Fisher Scientific Inc.). The mass spectrometer was operated in the data-dependent mode, in which a full-scan MS (from m/z 375 to 1,200 with the resolution of 60,000) was followed by MS/MS of the 30 most intense ions [15,000 resolution; normalized collision energy - 30%; normalized automatic gain control target (AGC) – 50%, maximum injection time - 30 ms; 60 sec exclusion].

#### LC-MS data processing

The raw files were searched directly against the human database updated in 2025 with no redundant entries, using Byonic (Protein Metrics) and Sequest search engines loaded into Proteome Discoverer 3.1 software (Thermo Fisher Scientific Inc., RRID:SCR_014477). The samples were searched in four batches and then compiled together into a single file. MS1 precursor mass tolerance was set at 10 ppm, and MS2 tolerance was set at 20 ppm. Search criteria included a static carbamidomethylation of cysteines (+57.0214 Da), variable modifications of oxidation (+15.9949 Da) on methionine residues and acetylation (+42.011 Da) at N terminus of proteins. Search was performed with full trypsin/P digestion and allowed a maximum of two missed cleavages on the peptides analyzed from the sequence database. The false-discovery rates of proteins and peptides were set at 0.01. All protein and peptide identifications were grouped, and any redundant entries were removed. Only unique peptides and unique master proteins were reported.

#### Label-free Quantitation Analysis

All data were quantified using the label-free quantitation node of Precursor Ions Quantifier through the Proteome Discoverer v3.1 (Thermo Fisher Scientific Inc., RRID:SCR_014477). For the quantification of proteomic data, the intensities of peptides were extracted with initial precursor mass tolerance set at 10 ppm, minimum number of isotope peaks as 2, maximum ΔRT of isotope pattern multiplets – 0.2 min, PSM confidence FDR of 0.01, with hypothesis test of ANOVA, maximum RT shift of 5 min, pairwise ratio-based ratio calculation, and 100 as the maximum allowed fold change. For calculations of fold-change between the groups of proteins, total protein abundance values were added together, and the ratios of these sums were used to compare proteins within different samples.

#### Isolation of peripheral blood mononuclear cells

PBMCs were isolated from commercially available freshly-collected buffy coat from healthy individuals (SER-BC-SDS, Zen-Bio Inc.) using Lymphopure™ kit (426202, BioLegend Inc.) as per manufacturer instructions. Briefly, the buffy coat was diluted with an equal volume of 1X PBS (SH30256FS, Fisher Scientific Inc.). The diluted buffy coat was mixed with half volume of Lymphopure™ and centrifuged at 800xg for 20 min at room temperature without braking.

Mononuclear cell layer at the plasma: Lymphopure™ interface was collected without disturbing the other layers. The mononuclear cells were washed in RPMI culture medium supplemented with 10% FBS and 1X penicillin/streptomycin and centrifuged at 500xg for 10 min at 4 °C. The cell pellet was resuspended in 100% FBS and the total number of cells was counted using a hemocytometer (02-671-54, Fisher Scientific Inc.). The cells were either directly used for subsequent experiments or aliquoted in cryovials at 5 x 10^7^ cells per vial in 90% FBS supplemented with 10% DMSO, and cryopreserved. The cryovials were thawed as needed for subsequent experiments.

#### T cell proliferation and activation assay

Freshly isolated or cryopreserved PBMCs were cultured in complete RPMI overnight followed by labeling with eFluor™ 450 cell proliferation dye (65-0842-85, Thermo Fisher Scientific Inc.). To induce T cell stimulation, labeled PBMCs were seeded at 1 x 10^5^ cells per well in a 96-well plate precoated with anti-human CD3 antibody at 0.5 µg/mL (BioLegend Cat# 317302, RRID:AB_571927). The seeded cells were cultured in complete RPMI supplemented with 0.1% β-mercaptoethanol and soluble anti-human CD28 antibody at 0.5 µg/mL (BioLegend Cat# 302902, RRID:AB_314304) for 96 hr. The EV treatments or control (equivalent volume of PBS) were also included at the indicated doses. T cell proliferation was confirmed by observing cell number increases in control wells under the microscope. After 96 hr, cells were collected and stained with fluorophore-labeled anti-human CD3-PerCP (BioLegend Cat# 344813, RRID:AB_10641841), CD4-BV650 (BioLegend Cat# 344691, RRID:AB_2936743), and CD8-SV538 (BioLegend Cat# 344785, RRID:AB_3097328) antibodies. The percentage of proliferating CD3+, CD3+CD4+, and CD3+CD8+T cells was further assessed by determining the dilution of eFluor™ 450 signal by flow cytometry. Unstained and single-stained controls were included to ensure proper gating strategy (**Fig. S6a**).

#### Animals

NSG-DKO mice, also known as NOD.Cg-*Prkdc^scid^ H2-K1^b-tm1Bpe^ H2-Ab1^g^*^7^*^-em1Mvw^ H2-D1^b-tm1Bpe^ Il2rg^tm1Wjl^*/SzJ (RRID:IMSR_JAX:025216), were obtained from the Biological Evaluation Shared Resource Facility at Purdue University. Mice were maintained under pathogen-free and temperature- and humidity-controlled conditions on a 12-hr light/12-hr dark cycle and received normal chow. Mice that exhibited inadequate human immune system development, lost 20% of their initial body weight, or had subcutaneous tumors that were ulcerated or bigger than 1,000 mm^3^ were euthanized and excluded from the study.

#### Humanization of mice

The NSG-DKO mice were humanized as described here^27^. Briefly, newborn pups (day 2-4) were irradiated at a sublethal dose (∼3Gy/300rad) followed by intrahepatic injection of 1 x 10^5^ CD34+CD38- human hematopoietic stem cells (HSCs). At 6-7 weeks after HSC inoculation, peripheral reconstitution of the human immune system was monitored by flow cytometric evaluation of the blood of humanized mice for the presence of CD45+ immune cells and other major subsets – T lymphocytes (CD3+) and B lymphocytes (CD19+) (see **Table S1** for antibody information). Blood from non-humanized mice was used as controls for comparison.

#### In vivo xenograft study

HCC827 cells (1 x 10^7^ cells per mouse) were subcutaneously implanted into the right hind flank of humanized NSG-DKO mice (n=6), and tumor growth was monitored. When tumor volumes reached an average of ∼150 mm^3^, mice were injected intratumorally with 50 µg of total H358 EVs (n=6) or equivalent volume PBS (control, n=6) every 3 days for a total of 8 doses. Tumor growth was monitored by measuring tumor volume at the indicated time points using a vernier caliper and the formula, V = (L x W^2^)/2, where V is tumor volume, W is tumor width, and L is tumor length. Body weight was also recorded for the duration of the study. After 21 days of EV treatment, mice were humanely euthanized using CO_2_ in accordance with institutional guidelines followed by cervical dislocation. Tumors and spleens were collected by surgical resection for subsequent analyses. Each tumor was divided into two equal halves, with one half processed for immunohistochemical staining. The other half of the tumor and the spleen were processed for flow cytometry-based immunoprofiling.

#### Immunohistochemical staining of tumors

Freshly harvested HCC827 tumors were immediately fixed in 10% neutral buffered formalin (6764240, Fisher Scientific Inc.) for 24-36 hr at room temperature followed by transfer to 70% ethanol. The tumor tissues were embedded in paraffin blocks, sectioned, deparaffinized, and rehydrated before hematoxylin and eosin (H&E) or antibody staining. H&E staining and immunohistochemical staining for CD3 (Agilent Cat# A0452, RRID:AB_2335677) and CD8 (Abcam Cat# ab209775, RRID:AB_2860566) were performed on histology sections of HCC827 tumors. Stained slides were digitized using the Aperio FL ScanScope system (Leica Biosystems Imaging, Inc.) at the Purdue Histology Core. The densities of CD3+ and CD8+ lymphocytes in the tumor were quantified using ImageScope (RRID:SCR_014311). The quality control cutoff for low staining intensity was determined by a board-certified pathologist using H&E-stained reference slides for the same specimens.

#### Flow cytometry-based immunoprofiling of tumors, spleen, and blood

Harvested tumors were minced in RPMI 1640 supplemented with 5% FBS, 1X penicillin/streptomycin, 50 μM β-mercaptoethanol (Gibco), 5LULml^−1^ DNase I (QIAGEN), 0.85LmgLml^−1^ collagenase V (collagenase from *Clostridium histolyticum*; Sigma-Aldrich), 1.25LmgLml^−1^ collagenase D (collagenase from *C. histolyticum*; Roche), and 1LmgLml^−1^ Dispase II (Gibco) and incubated for 30 min at 37L°C. The digested tissue was filtered using a 70 µm sterile cell strainer and cells were pelleted by centrifugation at 350xg for 5 min at 4 °C. Splenic cells were obtained by gentle pressure-dissociation of the spleen using 1X PBS buffer (SH30256FS, Fisher Scientific Inc.), and then passed through a 70 µm sterile cell strainer and pelleted by centrifugation.

The cell pellets from tumor, spleen, and blood samples were incubated in 10 mL 1X RBC (red blood cell) lysis buffer (00-4333-57, Thermo Fisher Scientific Inc.) for 5 min at room temperature to lyse the RBCs. The lysis was stopped by addition of 20 mL 1X PBS and the cells were pelleted by centrifugation. The cell pellet was resuspended in 2 mL of FACS buffer (0.5% bovine serum albumin and 2 mM EDTA in 1X PBS) and incubated for 15 min at 4L°C with human Fc receptor binding inhibitor (eBioscience, 14-9161-71) and stained with indicated anti-human antibody for 30 min at 4L°C (**Table S1**). Cells were then washed and analyzed by flow cytometry using BD LSRFortessa™ Cell Analyzer (BD Biosciences). Unstained control and single-staining or IgG isotype controls were performed to ensure proper gating strategy, which is described in **Fig. S6b** and **S7**. Data were collected and analyzed with a BD FACSDiva (v.9.0) and FCS Express™ version 5 (RRID:SCR_016431), respectively.

#### Bioinformatics analysis

The survival analysis was conducted using the UCSC Xena (RRID:SCR_018938)^28^. For pan-cancer survival correlation with TGOLN2 copy number, “TCGA Pan-Cancer (PANCAN)” study was selected with “TGOLN2 copy number” as the first variable. For cancer-specific survival correlation with TGOLN2 transcript, the following studies were selected – “TCGA Lung Adenocarcinoma (LUAD)”, “TCGA Lung Squamous Cell Carcinoma (LUSC)”, “TCGA Lower Grade Glioma (LGG)”, “TCGA Glioblastoma (GBM)”, and “TCGA Bladder Cancer (BLCA)”. For each study, “TGOLN2 gene expression” was selected as the first variable. For overall survival correlation with TGOLN2 protein, “Pediatric Brain Tumor Atlas: CBTTC” study (PBTA) was selected with “TGOLN2: protein abundance (Tandem mass tag)” as the first variable. For all TCGA survival analyses, “sample_type” was selected as the second variable. For the PBTA study, “Tissue Type (NCIT)” was selected as the second variable. Sample types with “null” for any of the two variables as well as sample types corresponding to “Solid Tissue Normal”, “Recurrent Tumor”, “Primary Blood Derived Cancer – Peripheral Blood”, “Additional - New Primary”, “Metastatic”, and “Additional Metastatic” were excluded from the analysis. Only the “primary tumor” sample type was retained for the analysis, which was a total of 10,226 samples for PANCAN, 1002 samples for LUAD, 496 samples for LUSC, 528 samples for LGG, 166 samples for GBM, 406 samples for BLCA, and 195 samples for PBTA study.

The immune cell correlation analysis was conducted as described in the R script in the supplementary data, adopted from the TCGAplot R package^29^.

#### Statistics and reproducibility

Statistical analysis was performed using GraphPad Prism v10 (GraphPad Software, LLC, RRID:SCR_002798). The number of replicates for each experiment is indicated in the corresponding figure legends. For western blotting data, raw blots are provided in the supplementary data. The two-tailed t-test with Welch correction was used to compare two groups and one-way ANOVA test with Tukey correction to compare more than two groups. Because the study was focused on NSCLC that affects both men and women, mice from both sexes were included. Statistically significant *p*-values are as indicated in the corresponding figure legends.

## Data availability

All source data generated during this study are provided with this paper and its supplementary data files. Any additional information is available from the corresponding author upon a reasonable request.

## Code availability

Source code for immune cell correlation analysis is available at GitHub [Link].

## Acknowledgements

This work was supported by R01 CA226259 (ALK) and R01 CA205420 (ALK) and an American Lung Association Innovator Award INALA2023, IA-1059916 (ALK). The humanization of mice and monitoring of tumor growth were conducted by Biological Evaluation Shared Resource Facility supported by P30 CA023168. The authors acknowledge the assistance of the Purdue University Histology Research Laboratory, a core facility of the NIH-funded Indiana Clinical and Translational Science Institute. Electron microscopy was performed using instrumentation in the Purdue Electron Microscopy Center (RRID:SCR_022687).

## Notes

### Competing Interest Statement

The authors have declared no competing interest.

