## Supplemental Information for "Exosome-like biogenesis from the Golgi releases extracellular vesicles lacking conventional tetraspanins that mediate immune evasion in cancer"

The authors declare no potential conflicts of interest

**Figure S1**

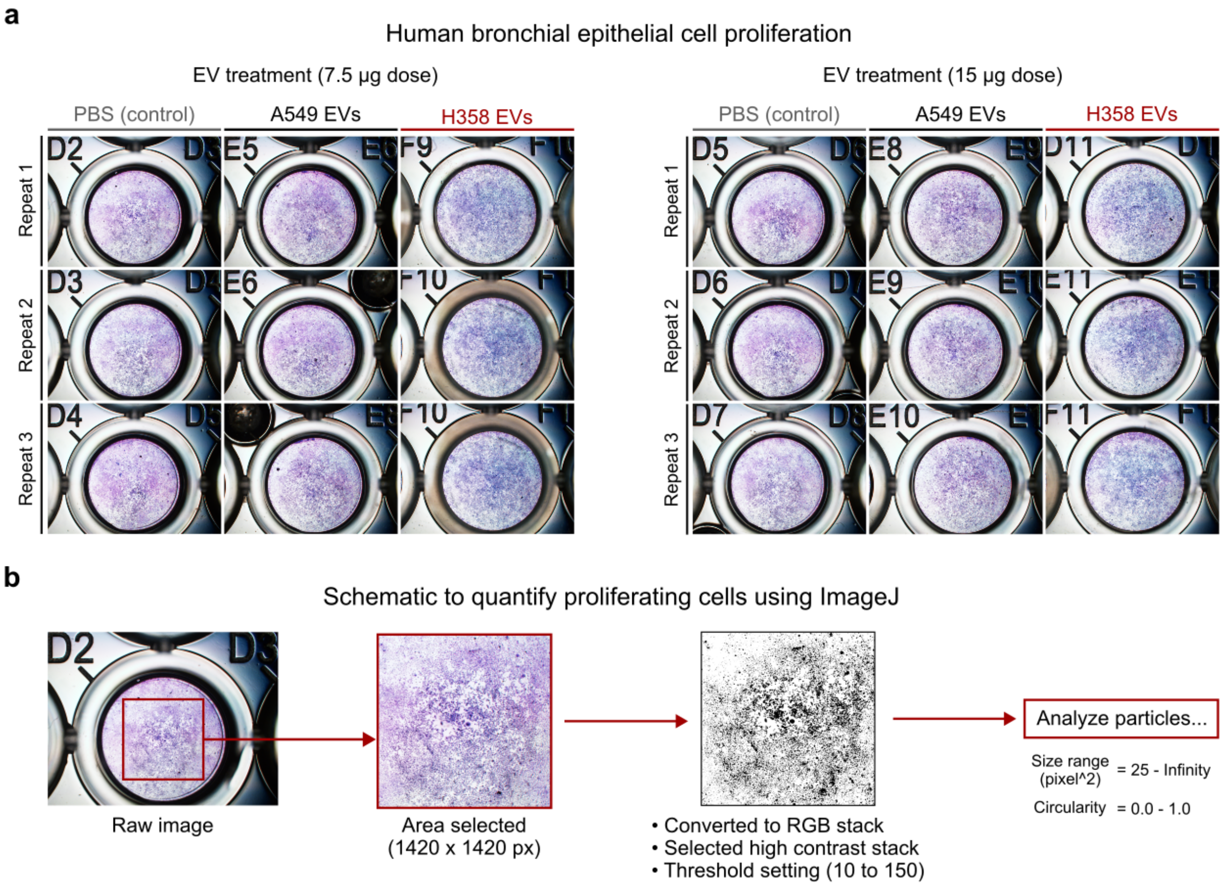


**Figure S1. Human bronchial epithelial cell proliferation.**

**a.** Raw images of human bronchial epithelial cells treated with A549 or H358 EVs (7.5 and 15 µg; 96 hr) compared with PBS control. **b.** Schematic to quantify the number of cells from each raw image using ImageJ.

**Figure S2**

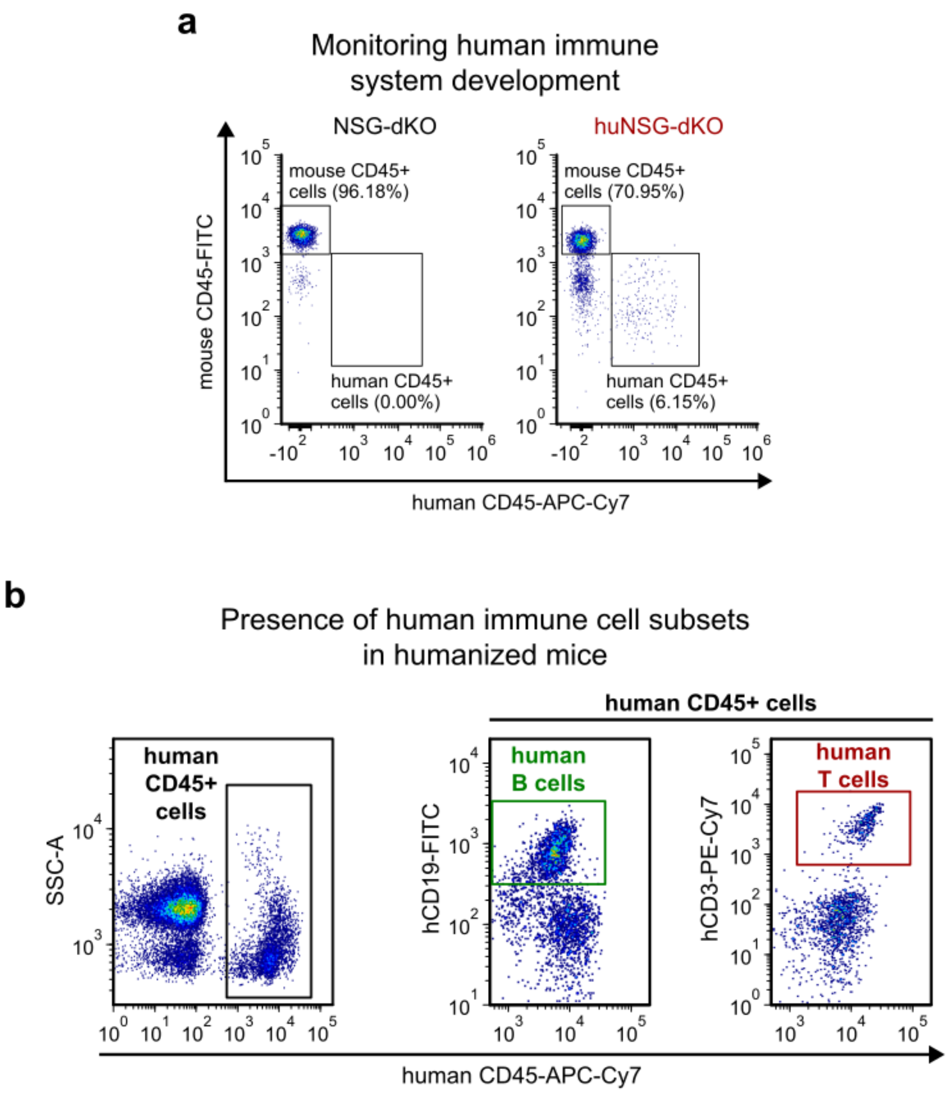


**Figure S2. Monitoring of human immune system development in huNSG-DKO mice.**

**a.** Representative dot plots quantifying presence of mouse and human CD45+ cells in peripheral blood of non-humanized (NSG-DKO) and humanized (huNSG-DKO) mice at 8 weeks of age. b. Representative dot plots depicting human CD45+ cells and human immune cell subsets – CD3+ T cells and CD19+ B cells – in peripheral blood of huNSG-DKO mice at 8 weeks of age.

**Figure S3**

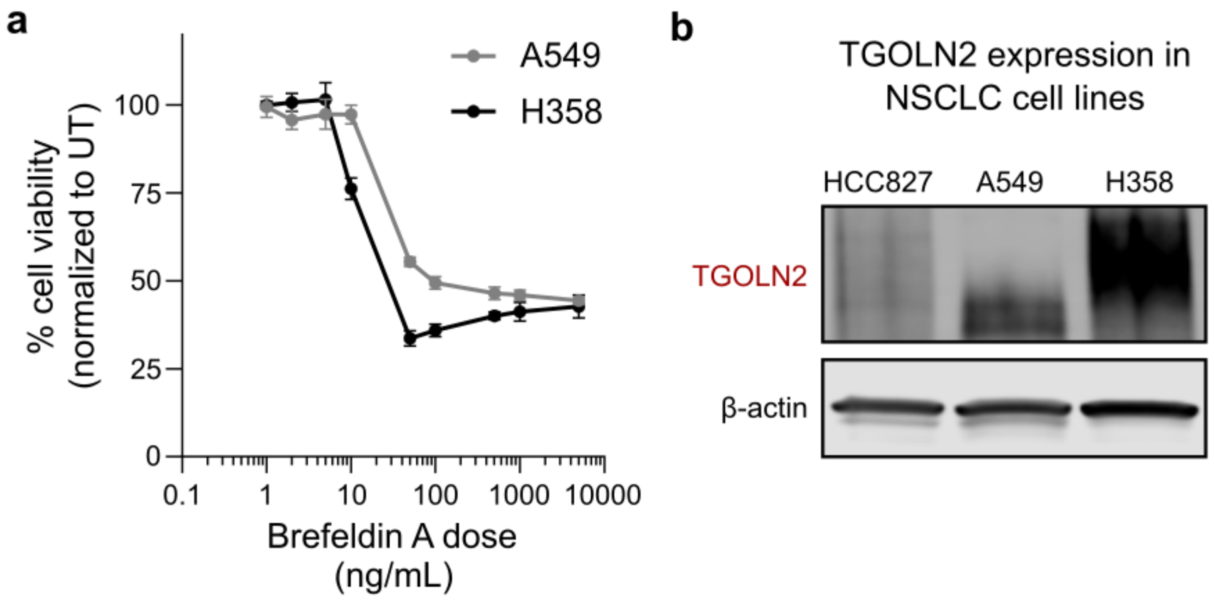


**Figure S3. Brefeldin A toxicity and TGOLN2 expression.**

**a.** Percentage viability of A549 and H358 cells after treatment with Brefeldin A for 48 hr at indicated doses (n=6). **b.** Representative immunoblot showing TGOLN2 expression in HCC827, A549 and H358 cells. β-actin is used as an endogenous control.

### Figure S4

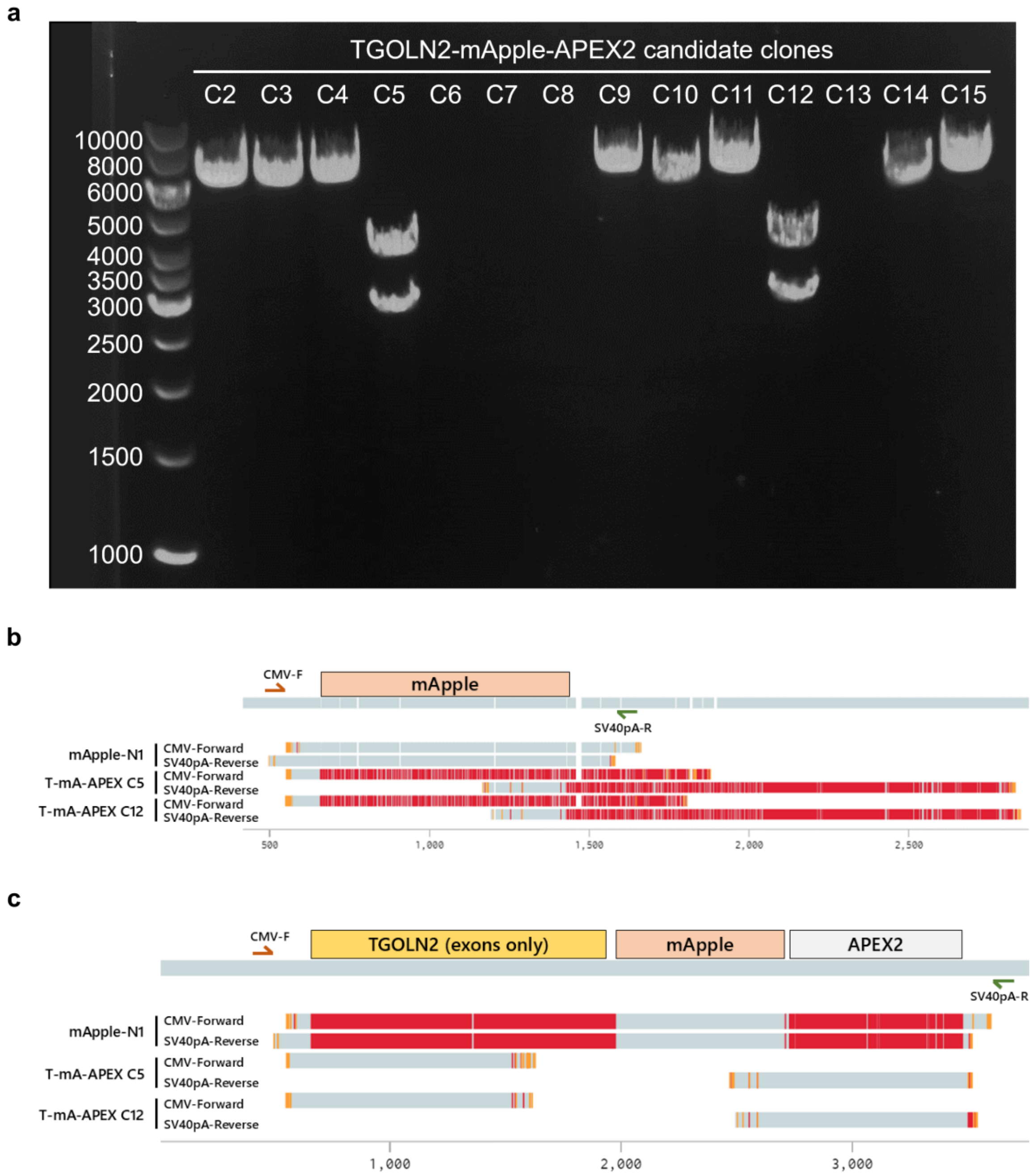


**Figure S4. Successful cloning of TGOLN2-mApple-APEX2 (TAPEX).**

a. Representative gel image showing *KpnI* and *MfeI* restriction digestion of candidate bacterial clones. Clones C5 and C12 show successful cloning of TAPEX sequence. Sanger sequencing was performed on mApple, TAPEX C5, and TAPEX C12 plasmids using CMV sequence as the forward primer and SV40pA sequence as the reverse primer. The sequencing results were aligned to the full *in silico* sequence of **b.** mApple plasmid or **c.** TAPEX plasmid. Red indicates no alignment and light blue indicates perfect alignment.

**Figure S5**

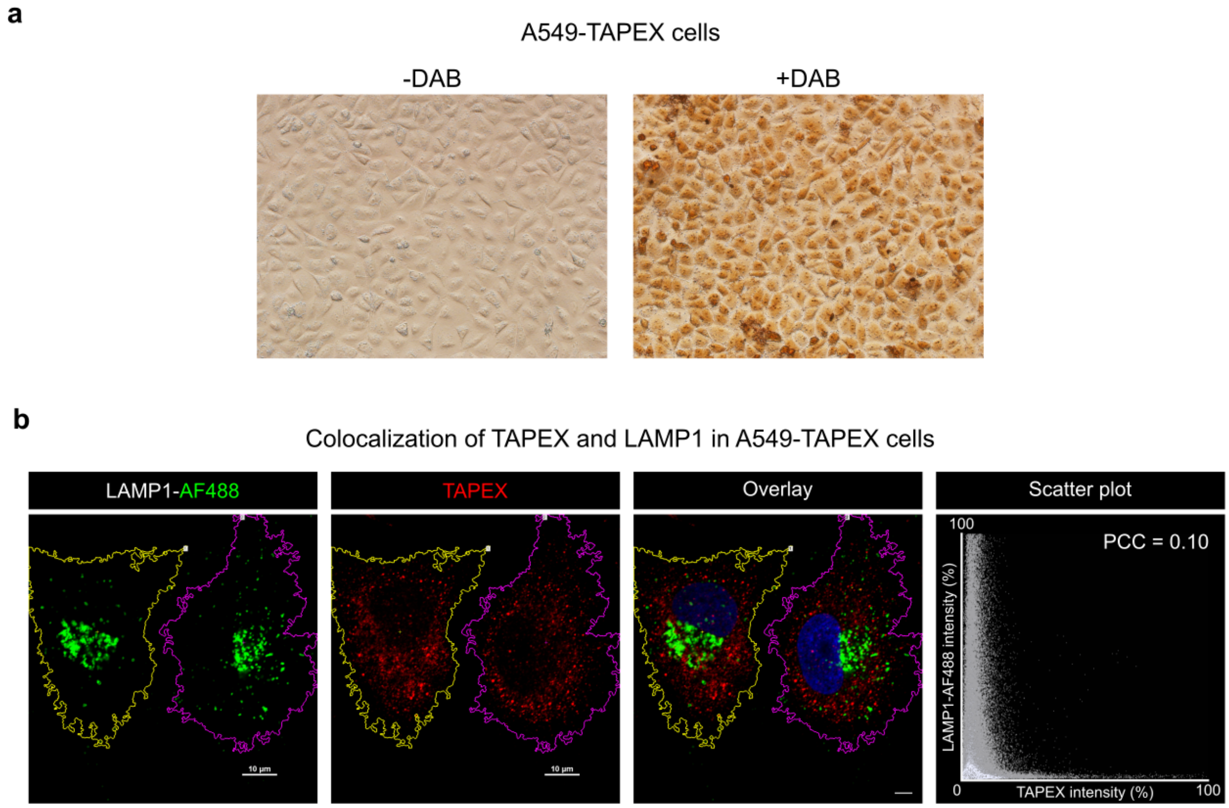


**Figure S5. A549-TAPEX validation and colocalization analysis.**

**a.** Bright-field imaging following 24 hr incubation of A549-TAPEX cells in the absence or presence of 1X Diaminobenzidine (DAB) solution with $H_2O_2$ (0.5 mg/ml DAB + 10 mM $H_2O_2$). **b.** Representative confocal imaging of A549-TAPEX cells showing distribution of the fluorescence signal for late endosomes (immunostained for LAMP1) and TAPEX (endogenous signal from mApply fluorophore). Hoechst is used for nuclear staining. Size bar – 10 µm.

**Figure S6**

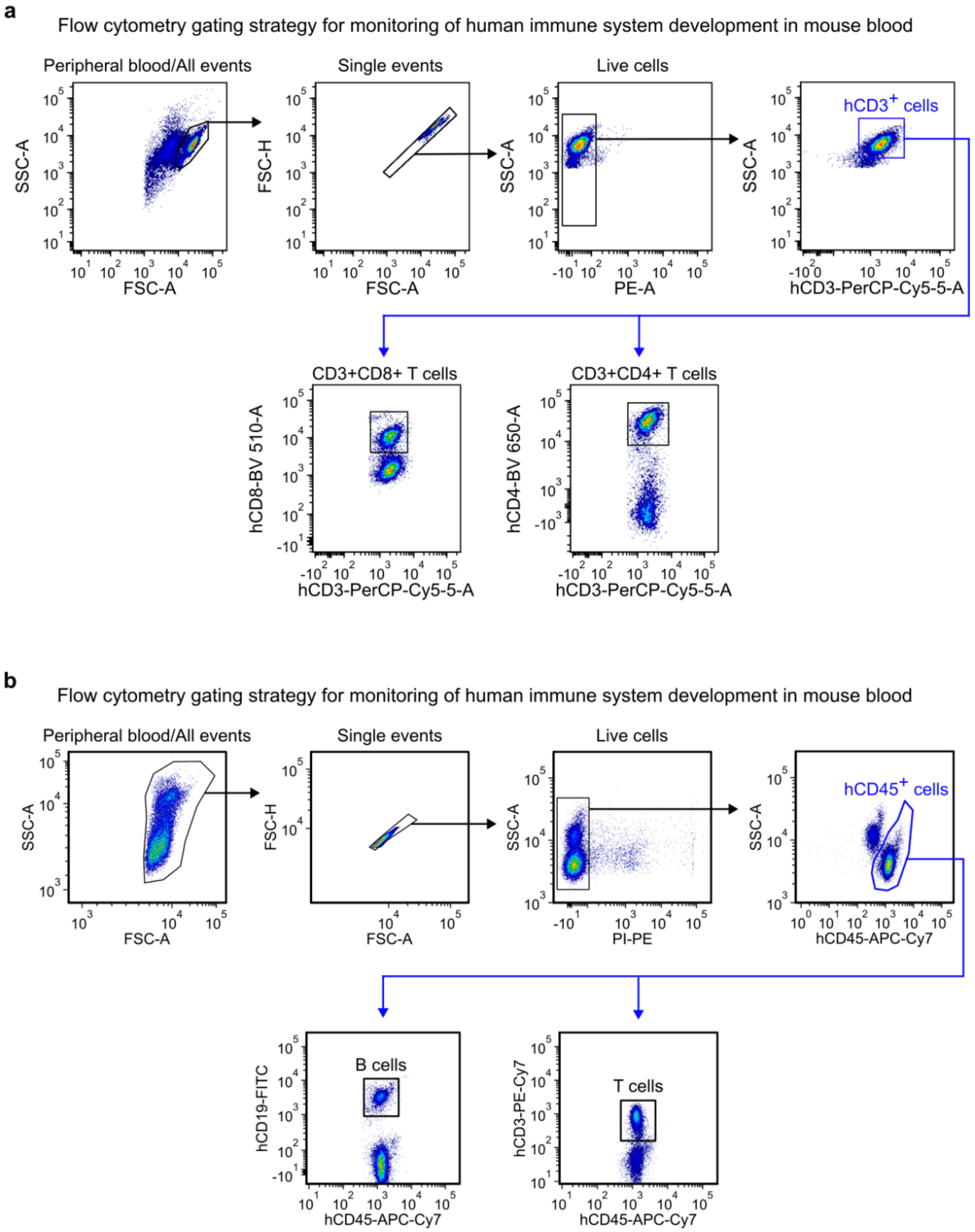


**Figure S6. Flow cytometry gating strategy for human primary T cell proliferation assay and monitoring human immune system development in peripheral blood.**

**a.** Gating strategy shown using untreated human primary T cells. **b.** Gating strategy shown using a representative blood sample from huNSG-DKO mice.

**Figure S7**

**a**

Flow cytometry gating strategy for immunoprofiling (representative tumor sample)

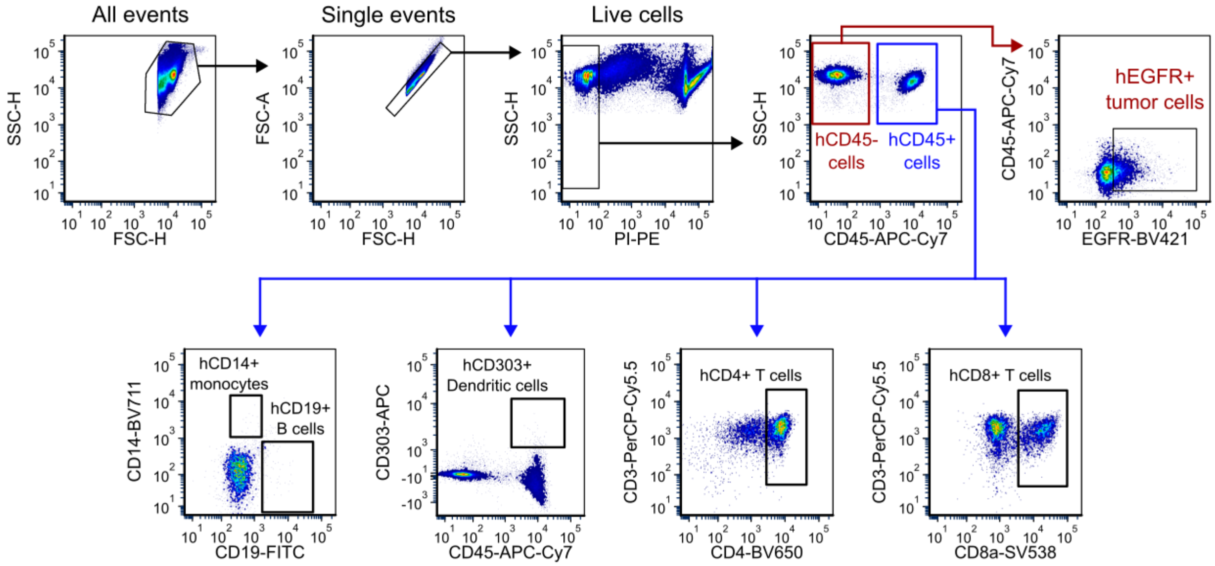


**b**

Flow cytometry gating strategy for immunoprofiling (representative spleen sample)

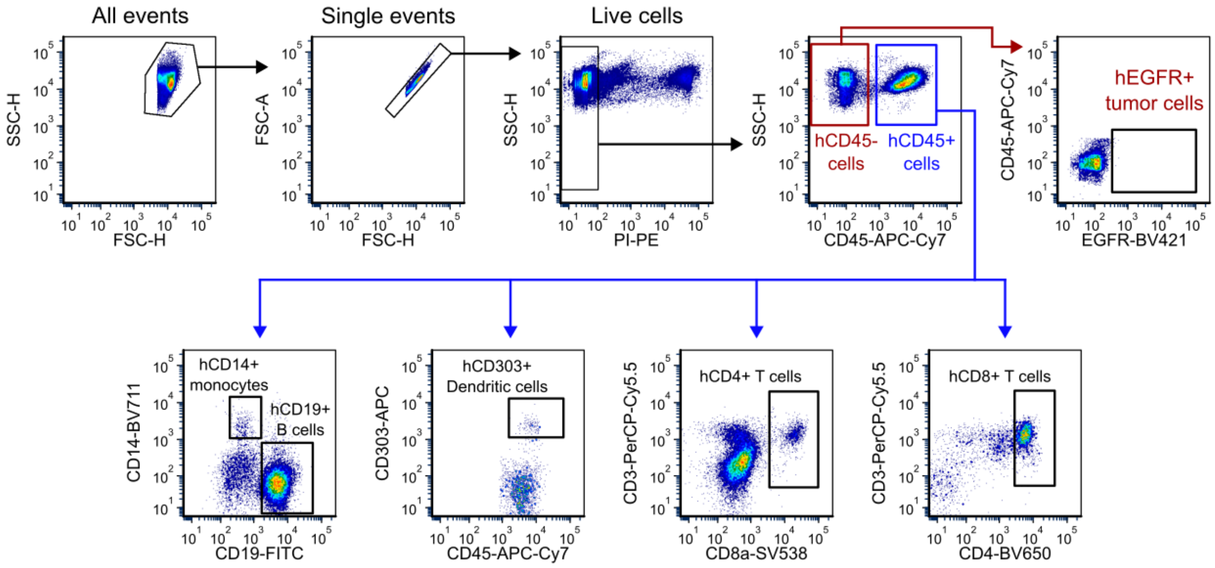


**Figure S7. Flow cytometry gating strategy for immunoprofiling of HCC827 tumors and spleen.**

**a.** Gating strategy shown using a representative HCC827 tumor from huNSG-DKO mice. **b.** Gating strategy shown using a representative spleen from huNSG-DKO mice.

**Table S1.** List of antibodies used in studies for western blotting, immunoprecipitation, flow cytometry.

| Primary antibody | Host/Target | Application | Manufacturer | Catalog # |
|---|---|---|---|---|
| CD45-APC-Cy7 | Mouse anti-human | Flow cytometry | Biolegend | 368516 |
| CD3-PerCP-Cy5.5 | Mouse anti-human | Flow cytometry | Biolegend | 317336 |
| CD4-Brillian Violet 650 | Mouse anti-human | Flow cytometry | Biolegend | 344692 |
| CD8a-Spark Violet 538 | Mouse anti-human | Flow cytometry | Biolegend | 388812 |
| CD19-FITC | Mouse anti-human | Flow cytometry | Biolegend | 302206 |
| CD303-APC | Mouse anti-human | Flow cytometry | Biolegend | 314407 |
| CD14-Brilliant Violet 711 | Mouse anti-human | Flow cytometry | Biolegend | 367139 |
| CD68-Brilliant Violet 785 | Mouse anti-human | Flow cytometry | Biolegend | 333825 |
| EGFR-Brilliant Violet 421 | Mouse anti-human | Flow cytometry | Biolegend | 352911 |
| CD16 | Mouse anti-human | Flow cytometry | Biolegend | 325902 |
| CD45-FITC | Rat anti-mouse | Flow cytometry | Biolegend | 103107 |
| CD3-PE-Cy7 | Mouse anti-human | Flow cytometry | Biolegend | 344815 |
| CD69-APC | Mouse anti-human | Flow cytometry | Biolegend | 310910 |
| TGOLN2 (TGN46) | Rabbit anti-human | Western blotting; immunoprecipitation | Fisher Scientific | NBP149643 |
| CD81 | Mouse anti-human | Western blotting; immunoprecipitation | ThermoFisher | 10630D |
| CD9 | Rabbit anti-human | Western blotting | Cell Signaling | 13174S |
| CD63 | Mouse anti-human | Western blotting; immunoprecipitation | Fisher Scientific | NBP232830F |
| IgG isotope control | Rabbit | Immunoprecipitation | Cell Signaling | 3900S |
| CD3 | Mouse anti-human | Primary T cell culture | Biolegend | 317301 |
| CD28 | Mouse anti-human | Primary T cell culture | Biolegend | 302901 |
